# Structural Analysis of 70S Ribosomes by Cross-Linking/Mass Spectrometry Reveals Conformational Plasticity

**DOI:** 10.1101/2020.05.04.077503

**Authors:** Christian Tüting, Claudio Iacobucci, Christian H. Ihling, Panagiotis L. Kastritis, Andrea Sinz

## Abstract

The ribosome is not only a highly complex molecular machine that executes translation according to the central dogma of molecular biology, but also an exceptional specimen for testing and optimizing cross-linking/mass spectrometry (XL-MS) workflows. Due to its high abundance, ribosomal proteins are frequently identified in proteome-wide XL-MS studies of cells or cell extracts. Here, we performed in-depth cross-linking of the *E. coli* ribosome using the amine-reactive cross-linker diacetyl dibutyric urea (DSAU). We analyzed 143 *E. coli* ribosomal structures, mapping a total of 10,771 intramolecular distances for 126 cross-link-pairs and 3,405 intermolecular distances for 97 protein pairs. Remarkably, 44% of intermolecular cross-links covered regions that have not been resolved in any high-resolution *E. coli* ribosome structure and point to a plasticity of cross-linked regions. We systematically characterized all cross-links and discovered flexible regions, conformational changes, and stoichiometric variations in bound ribosomal proteins, and ultimately remodeled 2,057 residues (15,794 atoms) in total. Our working model explains more than 95% of all cross-links, resulting in an optimized *E. coli* ribosome structure based on the cross-linking data obtained. Our study might serve as benchmark for conducting biochemical experiments on newly modeled protein regions, guided by XL-MS. Data are available via ProteomeXchange with identifier PXD018935.

## Introduction

Ribosomes, the molecular machines that are responsible for protein synthesis, have frequently attracted interest both from a biological [1] as well as from a methodological perspective [2–4]. Due to their high abundance in cells, they are straightforward to isolate [5], making them an ideal specimen for developing X-ray crystallography [6] and electron microscopy (EM) methods [2]. Ribosomes have been used in EM to develop positive and negative staining methods [7], evaluate the efficiency of direct electron detectors [8], and they have served as a model system to develop image processing protocols [2]. Due to their central role in protein translation, ribosomes possess an immense biological value, recognized by the award of the 2009 Chemistry Nobel Prize to Ramakrishnan, Steitz and Yonath “*for studies of the structure and function of the ribosome*”. Structural insights into ribosomes from various organisms, in a variety of conformations [9] and higher-order states [10, 11], in complex with other biomolecules [12, 13], inhibitors or antibiotics [14, 15] have been reported. These studies were influential to a detailed understanding of this highly intricate, MDa protein machinery [1, 16].

The *E. coli* ribosome has frequently been used in proteome-wide cross-linking/mass spectrometry methods (XL-MS) as a testbed for structural mapping of cross-linked peptides, and for determining the overall fit to the ribosome structure [17–19]. Indeed, as ribosomes are the biomolecular complexes with the highest abundance in the cell, most of the cross-linking reactions in cell extracts or *in vivo* are concentrated on the ribosome. Therefore, a large number of cross-linked peptides are available for evaluating cross-linking efficiency. As an example, 207 high-resolution atomic structures of *E. coli* ribosomes are currently depostited in the PDB database (date: February 1, 2020; resolution < 4.5; < 25 entities per biological unit). Suprisingly, only 87% of the ribosome’s protein content has been structurally characterized in these complexes, meaning that more than 1,000 residues are highly flexible and impossible to recover in poor or absent electron densities.

XL-MS would allow providing structural information for all regions of the ribosome, including ordered as well as flexible regions, resulting in a detailed understanding of the overall ribosomal architecture. XL-MS has matured from a method to study isolated biomolecules into a proteome-wide method to understand cellular protein interactions [20, 21]. Towards this goal, novel cross-linkers have been developed to optimize the discovery of cross-linked peptides with sensitive LC/MS/MS protocols [17, 19, 22, 23] and attemps have been made to estimate false discovery rates (FDR) for cross-links [24]. However, an important validation for proteome-wide XL-MS is the mapping of cross-links on structural models. There is still development in the field regarding the distance measure that is best applicable for mapping cross-links, i.e., Euclidian *versus* surface-exposed distance [25]. Also, the usefulness of FDR calculations and their correlation to true positive protein-protein interactions and the corresponding structural models [26] as well as the choice of molecular models that are used for cross-link mapping are in some cases suboptimal. This is because (a) current studies have a bias for high-abundant proteins, but methods to address this issue are being implemented [27] and (b) few molecular models deposited in structure databases are being evaluated for cross-linking distances, and only one molecular model is selected for distance calculation per protein complex, despite the wealth of structural data. This obvious limitation subsequently confines recovered results, which could be of relevance for the structural biology and function of the protein of interest.

Here, we apply the *N*-hydroxysuccinimide (NHS) ester diacetyl dibutyric urea (DSAU) as an amine-reactive, urea-based MS-cleavable cross-linker [28]. It possesses a spacer length of ~10 Å and is shorter than the widely-used cross-linker DSBU. Therefore, DSAU allows a distinct subset of distance constraints during structural mapping and/or subsequent modeling to be measured. We cross-linked the *E. coli* ribosome with DSAU and recovered 126 intra- and 97 intermolecular cross-links at an FDR of 1%. We then comprehensively mapped *all* identified cross-links onto *all* suitable *E. coli* ribosomal structures. Based on the satisfaction of those data, we eventually (a) remodeled flexible protein regions and discovered underlying conformational plasticity, (b) localized the ribosome-associated chaperone, the trigger factor, (c) completed protein structures with additional residues and domains, and (d) unveiled higher-order ribosome states. We finally highlight the broad synergy of XL-MS with high-resolution structural methods, as our XL-MS experiments allowed remodeling of 2,057 residues in total, optimizing the current working model of the *E. coli* ribosome (Workflow, **Fig. 1**).

**Figure 1.**
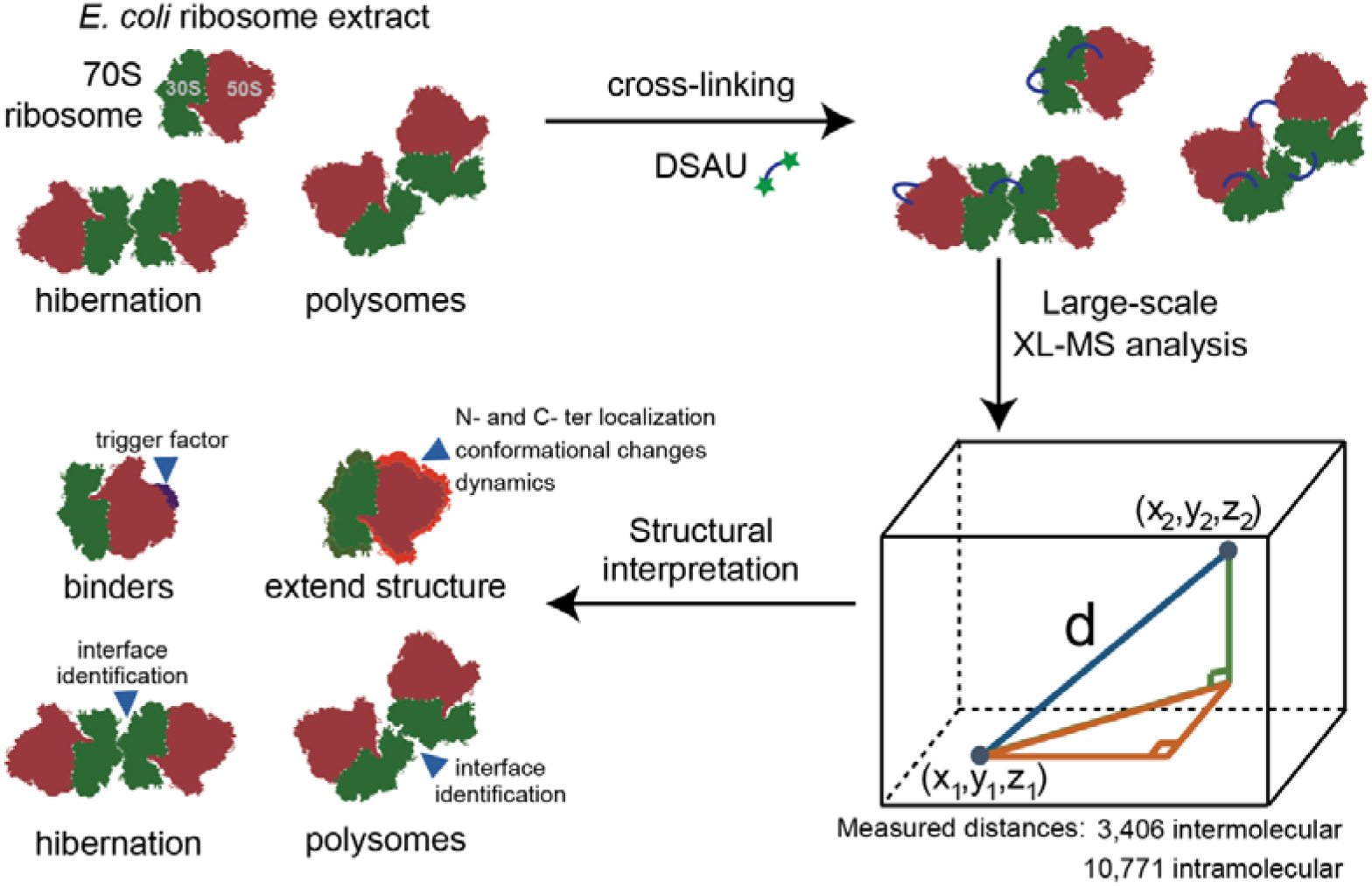
Workflow. We used the DSAU cross-linker with the 70S *E. coli* ribosome to identify binders, flexible regions, higher-order states and modeling of regions previously unresolved using derived data and structure-wide available data in the Protein Data Bank (https://www.rcsb.org/).

## Results

### Large-scale mapping of cross-links on *E. coli* ribosome structures

As mentioned above, we have systematically recovered 126 intramolecular cross-links and 97 intermolecular cross-links at 1% FDR, mapping 47 proteins and 36 protein-protein interactions in the *E. coli* ribosome (**Table S1**). We first analyzed the intramolecular cross-links. 126 non-redundant intramolecular cross-links were mapped onto 142 ribosome structures, while only 11% (14/126) of these cross-links could not be mapped to any available structure (**Table S2**). The cross-links show a distance distribution (*d*_intra_) that recapitulates the length of the DSAU cross-linker (N=10,769; median(*d*_intra_)=13.4 Å). As intramolecular ribosomal cross-links, except for L7/L12, do not involve protein-protein interactions, we set a threshold of 30 Å for evaluating those cross-links. By applying this threshold, 91.1% (102/112) of the mapped intramolecular cross-links were statisfied. As only intermolecular cross-links capture protein-protein-interactions in the ribosome, the intramolecular cross-links were not further analyzed, except for the interesting case of L7/L12, which is the only known multimeric protein in the *E. coli* ribosome.

For the 97 intermolecular cross-links, we identified 69 ribosomal structures with at least one intermolecular cross-linking pair (**Table S3**). The cross-links show a distance distribution (*d*_inter_) higher than that of intramolecular cross-links, clearly indicating flexibility in the respective ribosomal protein-protein interactions (N=3,304; median(*d*_inter_)=20.28 Å). Out of the 97 cross-links, 54 could be mapped on ribosomal structures, while 44 were novel (**Fig. 2A**), signifying that the residues involved in the cross-linking reaction are not represented in any published ribosome structure (< 3.5 Å resolution). We systematically evaluated the threshold of the DSAU cross-linker for distance measurements of the intermolecular cross-links detected (**Fig. S1**). Interestingly, the number of violated cross-links remains constant up to a threshold of 40 Å (**Fig. S1**). Therefore, we chose a distance threshold of 37.5 Å as this (a) accounts for the extensive conformational flexibility of the ribosome (it is 150% higher compared to the atomic distance of 25 Å which is the maximum Ca-Ca distance of cross-linked Lys residues, (**Fig. S2)** and (b) does not influence the recovery of higher false positive numbers (**Fig. S1**).

**Figure 2.**
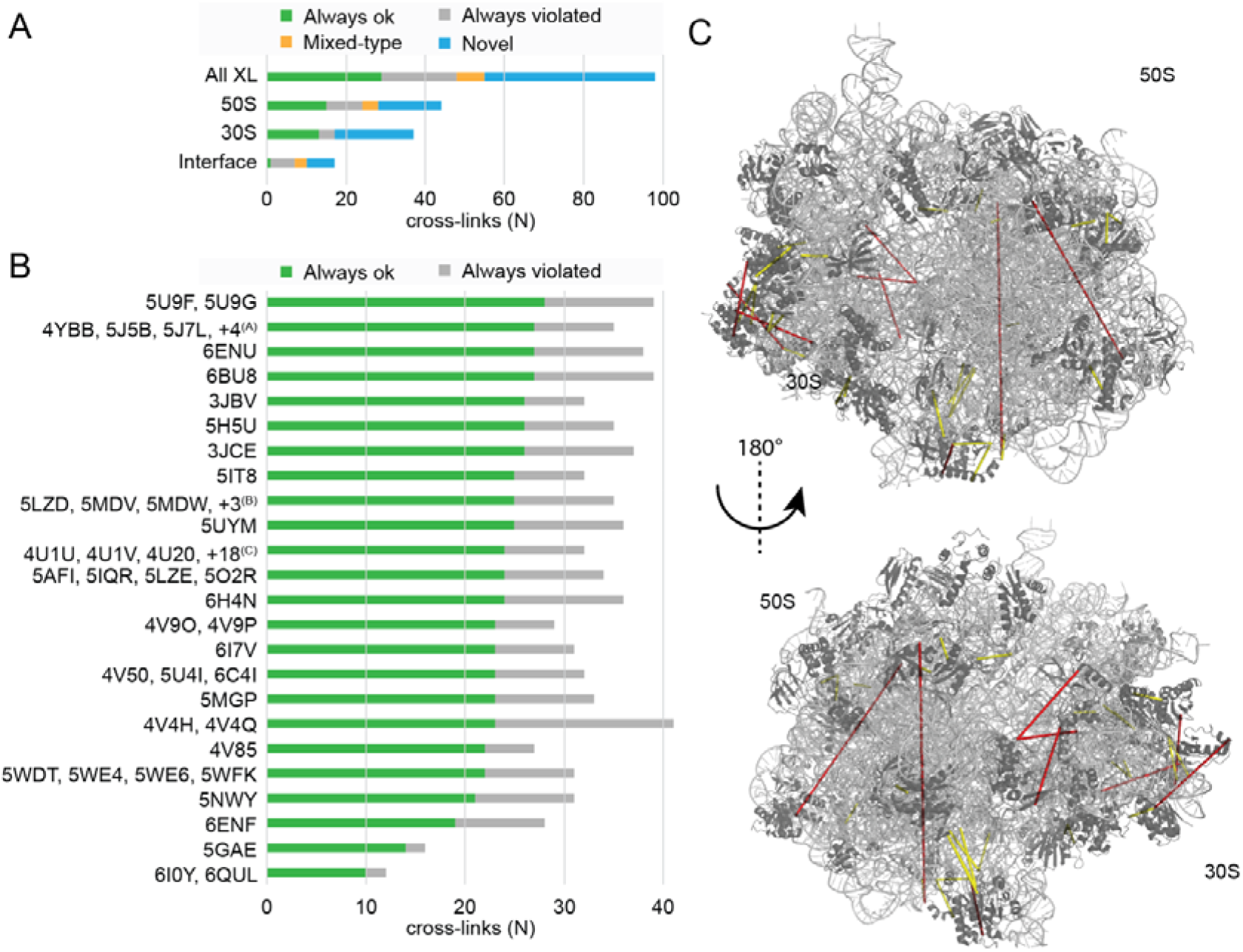
Statistics of database-wide mapping of ribosomal inter-molecular cross-links. (A) Classification of cross-links. Satisfied: cross-links always in the allowed range; Mixed-typed: cross-links both in the allowed and in the violated range, depending on the ribosome structure; Violated: cross-links always outside the allowed range; Novel: cross-links that could not be mapped on any high resolution ribosomal structure. (B) Classification of mappable cross-links on published structures. PDB IDs (www.rscb.org) with the same count of satisfied and violated cross-links are grouped. Pruned pdb-codes: ^(A)^ 5J88, 5J8A, 5J91, 5JC9; ^(B)^ 5MDY, 5MDZ, 6HRM; ^(C)^ 4U24, 4U25, 4U26, 4U27, 4V52, 4V54, 4V57, 4V64, 4V6C, 4V7S, 4V7T, 4V7U, 4V7V, 4V9C, 4V9D, 4WF1, 4WOI, 4WWW. (C) Cross-links on 5U9F, the published structure with the highest number of satisfied cross-links. Cross-links that satisfy the distance range are shown in yellow dashed lines; distances that are violated are shown in red. Ribosomal proteins (black) and rRNA (grey) are shown in cartoon presentation.

Therefore, 65% (35 out of 54) of the mapped cross-links are satisfied in at least one ribosomal structure (**Fig. 2B**). In particular, the ribosomal structure of the “ArfA-RF2 ribosome rescue complex” (5U9F) [29] has a wide coverage of satisfied cross-links, amounting for 28 non-redundant residue pairs in total (**Fig. 2B, 2C**). Eventually, we are able to explain, clarify, and satisfy 89% (17 out ouf 19) of the violated cross-links, and therefore, confidently map >95% of cross-links to the available ribosome structures.

### Plasticity of ribosomal active sites recapitulated by XL-MS

#### Translation initiation and codon probing

For efficient translation initiation, the 30S protein S21 is essential and consists of two α-helices, folding around the bound mRNA (PDB ID: 5U9F) [29]. Five cross-links were identified in S21, of which three map to the first α-helical region (residues 1-37) and two to the second (residues 38-71). Due to the localization of a violated cross-link (#86) in the second α-helix and the accessible space in the helical environment, only a planar movement of the second helical region can satisfy this cross-link. We performed a cross-link-driven reorientation of the second α-helix by molecular modeling, keeping the first α-helix rigid. By this, we could generate a model of S21, satisfying the violated cross-link in the second helix.

After initiation, aminoacyl-tRNA molecules must then be probed by trial and error, a procedure facilitated by the 50S ribosomal protein L7/L12, the only multimeric protein in the *E. coli* ribosome [30]. Due to its flexible nature, it is only partially resolved in few published structures (PDB IDs: 4V85 [31], 4V9O [32], 6I0Y [33]), however it was found to be highly cross-linked **(Fig. 3A)**. It is known that in *E. coli* up to four copies of L7/L12 can bind to a single ribosome [34]. We identified 15 novel interactions involving L7/L12 that include three different proteins, L6, L10 and L11 (**Fig. 3A**), while one intermolecular and 21 intramolecular cross-links are located in L7/L12 itself. In addition to cross-links which clearly indicate that L7/L12 is multimeric, the intramolecular cross-links may also reveal a higher-order oligomerization state. For this, we mapped the intramolecular cross-links on the published monomeric structure of the C-terminal (*C-ter*) domain of L7/L12 (PDB ID: 1CTF) [35] and measured both Euclidian and surface accessible distances using the xWalk software [36] (**Table S4**). Despite the Euclidian distances of all cross-links being in range (< 30 Å), the surface accessible distance for eight cross-links was above 30 Å, pointing to a higher-order state.

**Figure 3.**
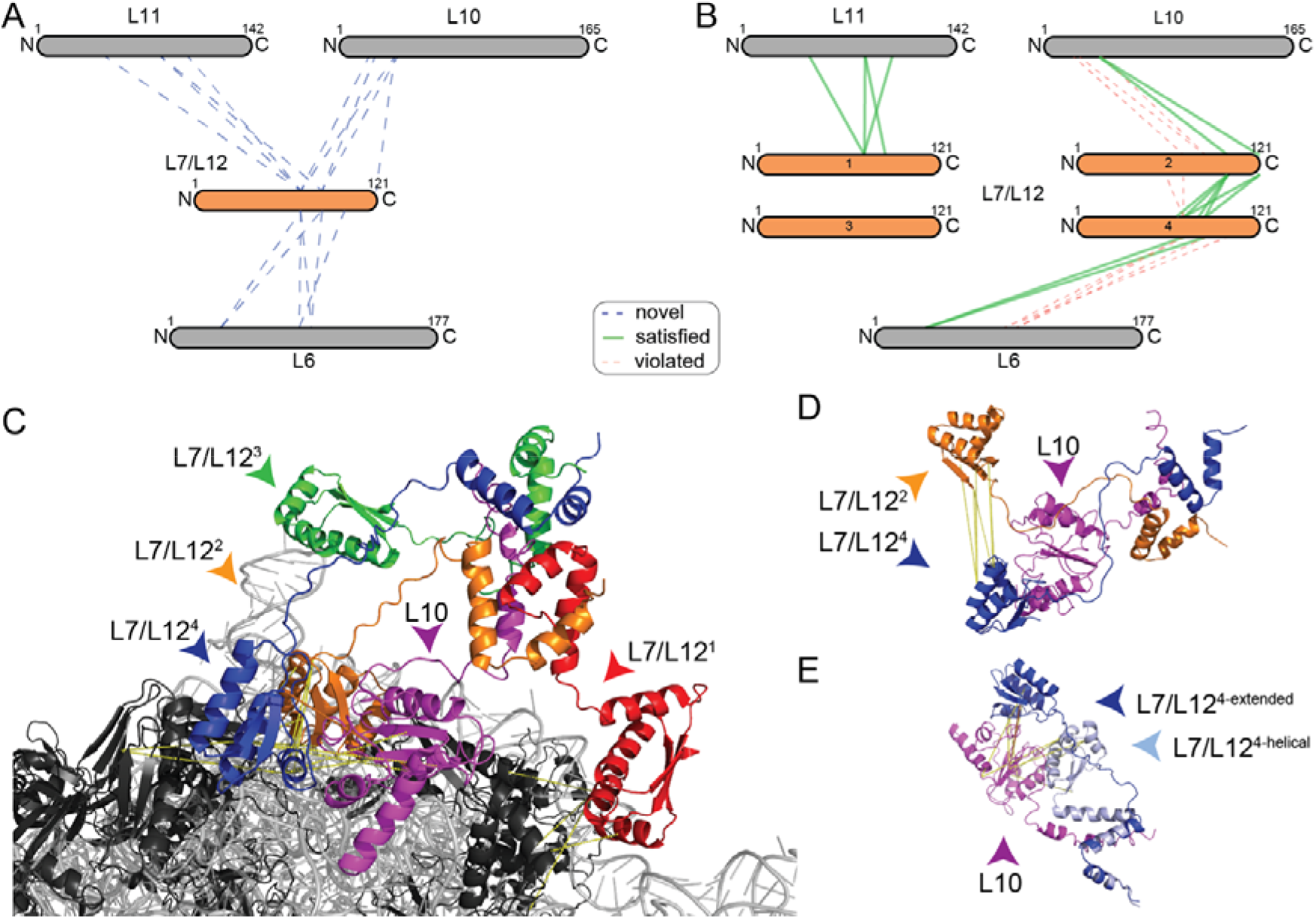
The dense interaction network of L7/L12 captured by XL-MS. Ribosomal proteins (black) and rRNA (grey) are shown in cartoon. Cross-linked proteins are color highlighted. Cross-links are shown as dotted lines, satisfaction (yellow) or minor violation (orange) are indicated. (A) Interaction network of L7/L12. Sizes of the boxes are scaled to the length of the protein sequences; each dotted line represents one identified novel cross-link pair. (B) Satisfied cross-links after remodeling; Only 4 cross-links of L6 to L7/L12 could not be unambiguously mapped (not shown); In addition, 5 intra-links show violation in the monomer, but satisfaction in the L7/L12 multimer. (C) Model of the interaction of an L7/L12 tetramer (blue, green, red, orange) bound to the helix 8 of L10 (magenta) based on the cross-linking information (see text). (D) Satisfaction of intra-molecular cross-links between CTD domains of L7/L12. (E) Inter-molecular cross-links satisfying both extended and helical conformation of the linker region of L7/L12; Note that all cross-links are satisfied in the helical linker conformation, and all but one cross-link are satisfied in the extended conformation.

Considering this information, we generated a tetrameric model of L7/L12 binding to the ribosome. We generated a tetramer because it is known that in *E. coli*, up to four copies of L7/L12 can bind to a single ribosome and was previously partially resolved in a tetrameric state [31]. Our extended model satisfies 60% of the novel cross-links to other proteins (**Fig. 3B-C**) and 6 out of 8 intermolecular cross-links between copies of itself (**Fig. 3D**). In this model, different degrees of flexibility of the linker are visible: For one of the L7/L12 units, the cross-link pattern could be satisfied by either an α-helical or an extended fold (**Fig. 3E**).

#### Frameshifting and ratchet movement

Ribosomal frameshifting is accompanied by a ratchet movement of the 30S and 50S subunit, while inhibited states of the ribosome can “freeze” either of those. This plasticity is captured by two cross-links in the interface between 30S and 50S subunits: L5 was cross-linked to S13 and S19 in a mixed-type manner (**Fig. 4A**). Interestingly, cryo-EM structures resolve an open conformation, as indicated by a cross-link minor violation (38 to 40 Å), while X-ray structures capture various states of the rachet movement. As an example, the crystal structure of the *E. coli* ribosome in complex with kasugamycin (PDB ID: 4V4H) [37] was resolved with two ribosomes in the asymmetric unit with a variability in the 50S-30S interface: We mapped the cross-links and discovered conflicting behavior of measured distances for each of those ribosomes (#62, 18.9 Å and 34.5 Å; #73, 24.2 Å and 41.2 Å). This indicates, that even in crystallized ribosomes, different conformational states could be existing, underlining, again, the high flexibility.

**Figure 4.**
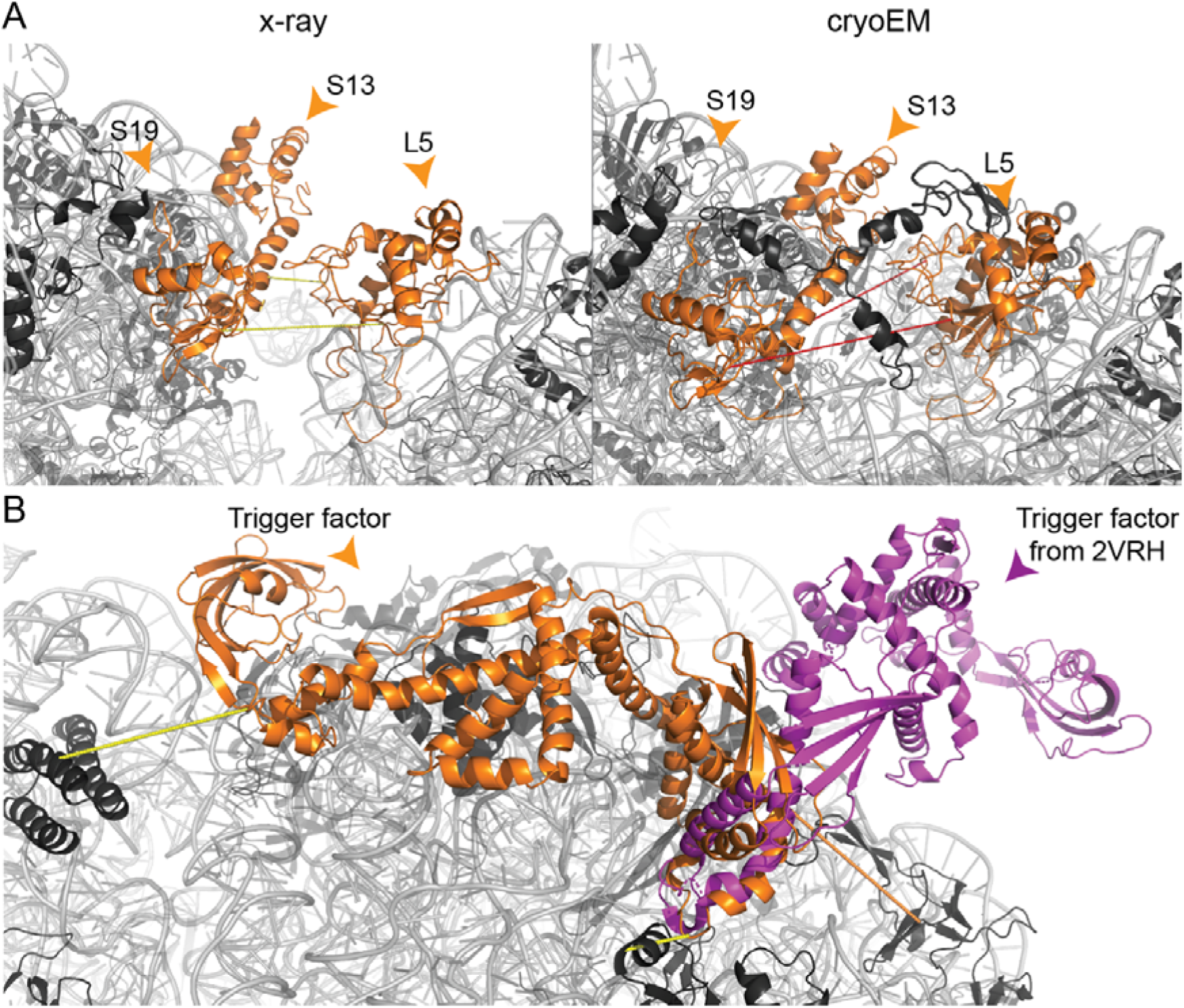
Visualization of differences in X-ray and cryoEM structures, identified by cross-links falling in the mixed-type category, and localization of the trigger factor. Ribosomal proteins (black) and rRNA (grey) are shown in cartoon. (A) Capturing the flexibility between the 30S and 50S subunits. In the X-ray structure (4V4H), cross-links #62 and #73 are satisfied (yellow dashed lines); in the cryoEM-structure (5U9F), a different conformation is visible, which violates the maximum distance (red dashed lines). (B) Structural model of the trigger factor (orange) in complex with the *E. coli* ribosome. Cross-link satisfaction is indicated as yellow lines, violation as orange lines. The known low-resolution structure (2VRH) is superimposed and shown in magenta.

#### Flexibility and rotation of the L1 stalk and unfolding of the mRNA by S3

The L1 stalk is a flexible substructure consisting of the ribosomal protein L1 and helices 76-78 of the 23S rRNA. We observed five cross-links from L1 to the L9 protein and one cross-link from L1 to the L33 protein. The available structures satisfy five out of six cross-links; the remaining cross-link (#1; 63.3 Å) reflects the known extensive conformational variability of the L1 stalk, including movements of up to 20 Å [38], which can be sampled only with extensive molecular dynamics simulations in combination with higher amount of data [39]. In addition to the L1 stalk movement, mRNA needs to be single-stranded during translation. The 50S protein S3 at the mRNA entrance tunnel is directly involved in the unfolding of the mRNA [40]. We identfied a set of five cross-links, four to S10 and one to S2, including the structurally unresolved 26-aa, C-terminal (*C-ter*) region of S3. We identified a unique structural homolog (PDB ID: 2JPL [41]; 38% similarity) using HHPRED [42] and modeled additional parts of the S3 protein. Without implementing any of the cross-linking constraints, the model satisfies all cross-links and, thereby, validates the structure with the additional *C-ter* region (**Fig. S3**).

#### Ribosomal chaperone

We identified three cross-links to the trigger factor, the only ribosome-associated chaperone in *E. coli* [43]. The bound state of the trigger factor is resolved at low-resolution (19 Å, PDB ID: 2VRH) [44] and shows an open conformation (**Fig. 4B**). It is also partially resolved in its *N*-terminal (*N-ter*) region in a homologous high-resolution ribosomal structure from *Haloarcula marismortui* (PDB ID: 1W2B [45]). In addition, the full-length structure in the unbound state has been solved by NMR spectroscopy (PDB ID: 1W26 [45]). Based on the published structures of the trigger factor homolog, a simple superposition of the unbound state on the bound *N-ter* region of the trigger factor resulted in a model that violated the cross-linking distances in the *C-ter* region, recapitulating the same binding mode observed in the low-resolution cryo-EM map previously published (PDB ID: 2VRH [44]). We therefore applied a cross-link-driven structural modeling of the trigger factor. We confirmed that the *N-ter* region is bound near the exit tunnel of the nascent peptide chain, formed by L23 and L24 [45]. The distance violation (50.3 Å) between the trigger factor and L24 is justified by the inherent flexibility of the exit tunnel. Interestingly, we discovered that the *C-ter* region folds back to the 23S RNA, forming an extensive non-covalent interaction network with the 23S-RNA surface (**Fig. 4B**).

### Extending the ribosomal model structure

We directly discovered that three cross-links involving the *N-ter* amine groups of S1, S8 and S18 could not be mapped on the ribosome because they were simply missing. By *de novo* completion, these novel cross-links could be mapped and were found to satisfy the distance threshold. However, the majority of missing residues are located in highly flexible regions and were subsequently identified and characterized, as described for L31 and S1 below.

#### Localization and C-terminal flexibility of L31

Based on the mixed-type distribution, meaning there is satisfaction in some, but violation in other structures, of a cross-link between L31 and L5, we were able to identify an annotation discrepancy for L31 in crystallographic structures of *E. coli* ribosome (PDB IDs: 4V4Q [46] and 4V4H [37]) (**Fig. 5A**). In the respective crystal structures, the electron density was mistakenly identified as L31 instead of L28. The averaged B-factor of L31 in both structures and both asymmetric units is remarkably high, 73.8 Å^2^ and 76.8 Å^2^ in 4V4Q and 77.8 Å^2^ and 80.3 Å^2^ in 4V4H [37], corroborating an inconclusive placement (**Fig. 5A**). In these structures, we also discovered 10 additional cross-links located in the *C-ter* region of L31 that are were greatly violated (above 150 Å) and result from the incorrect placement of L31. In addition, although L31 is correctly placed in recent ribosomal structures, the *C-ter* region is still unresolved, and therefore, cross-linking distances involving the *C-ter* cannot be mapped. In addition, the *C-ter* residues 63-70 (FNIPGSK) of L31 are known to be proteolytically cleaved by protease VII during purification [47]. However, we reveal that, by identifying these cross-links, L31 also exists in a non-cleaved state. This was confirmed by the identification of the *C-ter* peptide of full-length L31 during peptide mass fingerprint analysis (**Fig. S4**, **S5**). By *de novo* modeling of the additional residues at the *C-ter* of L31, we could explain four out of ten (40%) cross-links, indicating a high flexibility of the L31 *C-ter*. By using the cross-links as distance constraints in data-driven homology modeling, we were finally able to satisfy nine out of ten (90%) crosslinks. Conseqeuntly, experimental data suggest two distinct conformations of L31’s *C-ter* (**Fig. 5B**).

**Figure 5.**
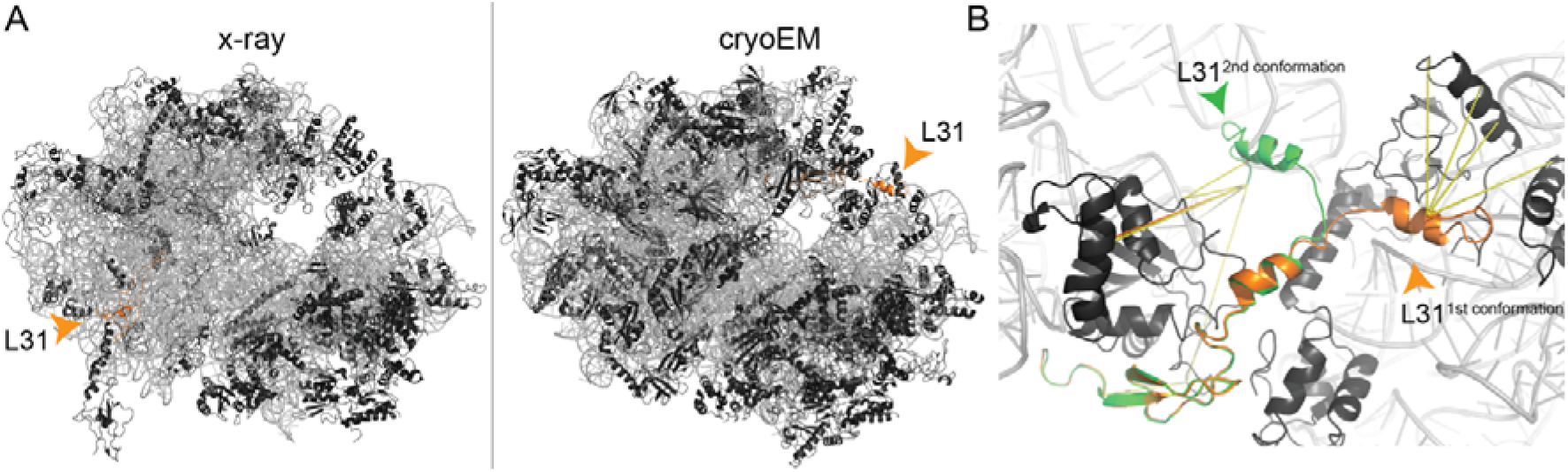
Localization and flexibility of the L31 ribosomal protein. Ribosomal proteins (black) and rRNA (grey) are shown in cartoon. L31, involved in cross-linking, is highlighted in orange. (A) Annotation discrepancy of L31 in the crystal structure (4V4H) in comparison to an recent cryoEM structure (5U9F). The orange arrow indicates the annotated localization at the ribosomal structure. (B) Two distinct conformations of the CTD of L31. In orange, the first conformation is shown, where cross-links #66, #71-72 and #74 are satisfied. In green, the second conformation is shown, where cross-links #27-30 and #39 are satisfied and cross-link #38 (orange line) is only minorly violated (38.8 Å). Cross-link #26, located in the NTD, is satisfied in both conformations.

#### Recapitulation of interaction networks for S1

A dense network of 18 cross-links (15 novel, two violated and one satisfied) between S1 and eight proximal proteins was identified in our XL-MS studies (**Fig. 6A**). S1 is the largest protein of the *E. coli* ribosome with a molecular weight of 61 kDa and is essential for docking and unfolding of structured mRNA [48], but for mRNAs with a strong Shine-Dalgarno sequence and short 5’-UTR, the S1 is not needed [48]. Only the *N-ter* domain (NTD), bound to the ribosome, was recently resolved in complex with the ribosome (PDB IDs: 6H4N, 6BU8) [49, 50]), and most of the other sequence regions of S1 are structurally uncharacterized (central domain and *C-ter* domain (CTD)). Biophysical studies suggest that the structure of S1 could be very elongated (up to 230 Å) [51], proposing a model of a bound NTD and a flexible CTD, which probes mRNA present in the cytosol. Here, we generated two conformations of the S1’s CTD that is represented by two bound states: For the structurally unknown central region (150-300), we identified a close structural homolog using HHPRED (*E. coli* elongation factor Ts; PDB ID: 4Q7J) [52]. In this region, five intermolecular cross-links were identified, of which two could be satisfied and one is only moderately violated (40.6 Å). The two additional cross-links are greatly violated (83.9 Å and 97.5 Å) and could thereby point to an extended conformation of S1 [51].

**Figure 6.**
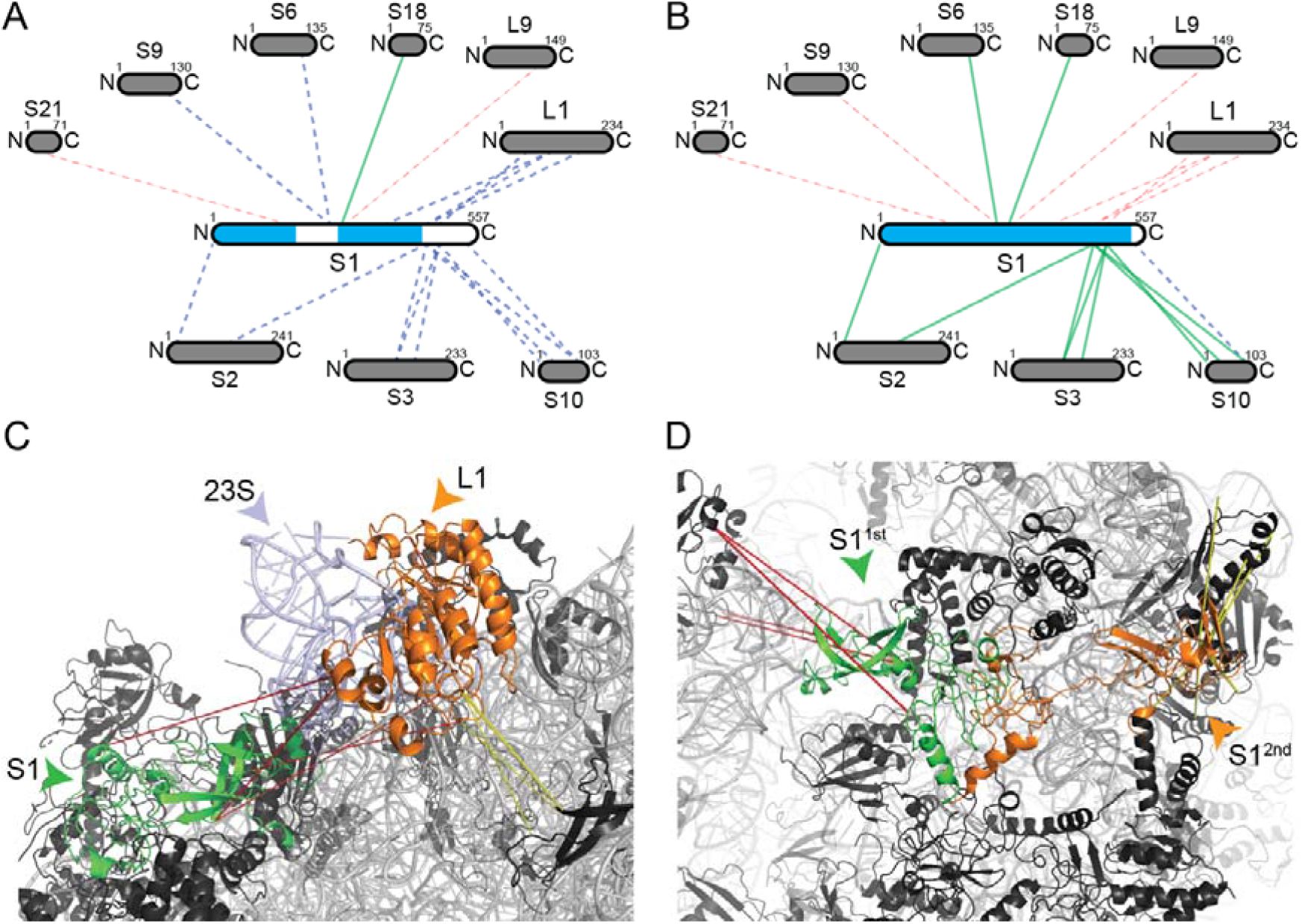
Identification and remodeling of the interaction network of S1. Interaction network of S1 before (A) and after (B) modeling. Sizes of the boxes are scaled to the length of the protein sequences and blue color in S1 indicates structurally characterized sequence parts. Each line represents one identified cross-link pair. Novel interactions are shown as dashed lines, satisfied (green) and violated (red) cross-link as colored lines. Ribosomal proteins (black) and rRNA (grey) are shown in cartoon. Remodeled proteins are highlighted in color (C-D). Cross-links are shown as dotted lines, and satisfaction (yellow) and violation (red) are also indicated. (C) Structural interaction network of the CTD of S1 (green) and the L1 stalk, formed by L1 (orange) and helices 76-78 of the 23S RNA (light blue). The cross-links between S1 and the L1 stalk show distance violation, but the L1 stalk itself to proteins in vicinity shows both, satisfaction (yellow) and violation (red) of cross-links, indicating different conformations as function of L1 stalk movement (see text for details). (D) Distinct conformations of the CTD of S1 (1^st^/2^nd^;green/orange). 1^st^ conformation (green) recapitulates the interaction of S1 with the L1 stalk in the 50S subunit. 2^nd^ conformation (orange) recapitulates an interaction with other 30S subunits and completely satisfies all cross-links.

Additionally, the majority of cross-links is located in the CTD of S1 that has so far been unresolved in existing ribosomal structures, but was resolved by solution NMR in its unbound state (free protein, PDB ID: 2KHJ) [53]. Based on these data, we modeled the structure of S1 in its bound state, where only the 30 last amino acids at the *C-ter* are missing (**Fig. 6B**, structural coverage indicated in blue). We discovered two states of the CTD: A highly complex interaction with the L1 stalk (see “*Flexibility and rotation of the L1 stalk”*), and a second interaction within the 30S subunit, as corroborated by four and seven cross-links. Our models show that either (a) the CTD domain of S1 folds on the 23S rRNA of the L1 stalk (**Fig. 6C**). where extensive conformational changes are predicted to occur [38] or (b) the CTD is bound to the 30S subunit that is validated by satisfying all seven cross-links (**Fig. 6D**).

### Multimeric states of ribosomes

In crystallographic ribosomal structures, two cross-links between L2 and L9 are simultaneously violated and satisfied as apparently, L9 undergoes a conformational change during crystallization. The extended state of L9 is found in 33 crystallographically determined ribosomal structures from *E. coli* (**Table S2**) and is imposed by crystallographic contacts (**Fig. 7A**) [9]. In ribosome hibernation, the dimeric interface is formed by the 30S subunit, while the 30S protein S2 of both ribosomes forms the core of the interface and 30S protein S1 is present in an inactive conformation [49]. The structure of the hibernated 100S ribosome has been solved by cryo-EM (PDB ID: 6H58) [49], but was not considered for mapping the cross-links due to its moderate resolution of 7.9 Å. Nevertheless, that structure is highly valuable in explaining two violated cross-links, which are now in good agreement with the low-resolution cryo-EM structure of the 100S ribosome, thereby further corroborating the low-resolution cryo-EM model (calculated distances 18.3 Å and 25 Å, **Fig. 7B**). Finally, distance measurements in polysomes are not feasible due to the absence of high-resolution structures as they are heterogeneous and form various assemblies [11]. Usually, ribosomes are connected “top-to-top” in polysomes, with a 30S-30S interaction along the mRNA body and the 50S subunits facing outwards in a pseudo-helical manner [11]. Cross-links spanning through the ribosomes might be explained by two adjacent ribosomes bound in a top-to-top manner. This has been observed for the two cross-links in the 50S with violated distances of 120.9 Å and 166.6 Å, when mapped on the monomeric ribosomal structure. It might well be possible that these cross-links are satisfied in higher-order polysomal states.

**Figure 7.**
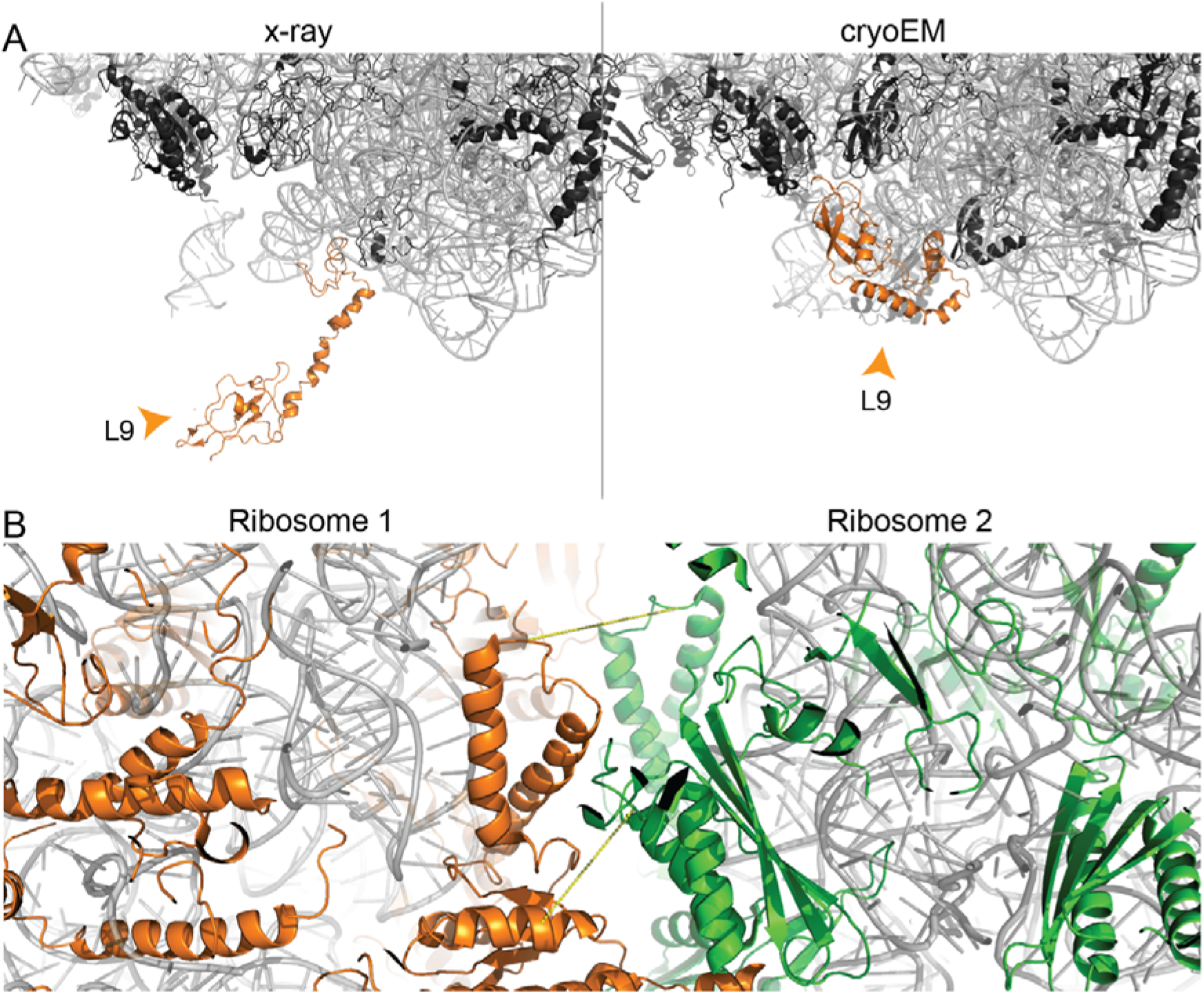
Multimeric states of the ribosome. Ribosomal proteins (black) and rRNA (grey) are shown in cartoon presentation. (A) Different conformations of L9. The protein is localized in the interface region of the asymmetric unit in the crystal structure (4V4H), and, thereby, is in an extended conformation. In the cryoEM structure (5U9F), the protein is folding back onto the surface of the ribosome. (B) Cross-linking reveals presence of hibernated ribosomes in the sample (PDB ID 6H58). Ribosomal proteins of the two ribosomes are color-highlighted (green/orange). Cross-links #81 and #97 are shown.

## Discussion

The published ribosomal structures of *E. coli*, albeit of impressively high numbers, account for a large fraction of ribosomal proteins, binders, cofactors, and translation states. Interestingly, our cross-linking experiments identified ~50% of cross-links that are novel, and are not part of published high-resolution published ribosomal structures. This means that current structural models only partially recapitulate the intrinsic flexibility of the 70S ribosome.

Our methodology, *i.e.* large-scale mapping of cross-linking data on *all* ribosomal structures and subsequent cross-link-based modeling, allowed us to capture novelties in technical, biochemical, and biological aspects. In particular, we have identified crystallographic contacts by visualizing violated cross-link distances, as in the case of L9. We have identified complex oligomerization states of L7/L12 and proposed a model for its structural role in the context of the ribosome; and we discovered discrete, but limited conformations for *N*- and *C-ter* regions of cross-linked ribosomal proteins S1 and L31. We have, ultimately, unraveled a novel interface for the *C-ter* region of the trigger factor, satisfying the cross-linking data. Higher resolution structures of trigger factor in complex with the *E. coli* ribosome are predicted to recapitulate our model where the trigger factor interacts with the 23S RNA in an extended conformation.

In our final *E. coli* ribosome model, we added or altered 2,057 residues (**Fig. 8**). In total, we were able to satisfy seven novel and four violated intramolecular, and 28 novel and 10 violated intermolecular cross-links, in addition to the already satisfied cross-links. This final model involves 115 intra-molecular cross-links and 71 inter-molecular cross-links, now fulfilling the distance threshold (30 Å for intra-, 37.5 Å for inter-molecular cross-links) with 91% and 73%, respectively **(Table S5)**. Our workflow of remodeling a single protein in a ridgid environment can be further improved by considering the environment of the protein to be flexible and suitable for remodeling, especially, if also rRNA is involved, as *e.g.* in the L1 stalk. Additionally, seven cross-links could not be satisfied for the L7/L12 multimer. Our proposed bound model of four L7/L12 proteins is a snapshot of the variable complexes that the multimer can adopt to probe aa-tRNAs, and therefore only a subset of cross-linking data is satisfied. The flexibility of the 30S-50S interface is further highlighted by five additional violated cross-links that are mapped on L5, S9 and S10, which are all known to be involved in tRNA binding [54] and frameshifting [55]. An exercise to finally fit all derived cross-links in a single static model or in snapshots is impossible, and highlights the fact that the ribosome, even in its single, purified state, probes a significantly large and complex conformational landscape. In addition, this poses several questions regarding the model if we systematically consider conformational ensembles of ribosomal proteins. This is because it is difficult to disentangle which constraint originates from which conformation. Nevertheless, our generated model is an averaged model, which incorporates multiple conformations.

**Figure 8.**
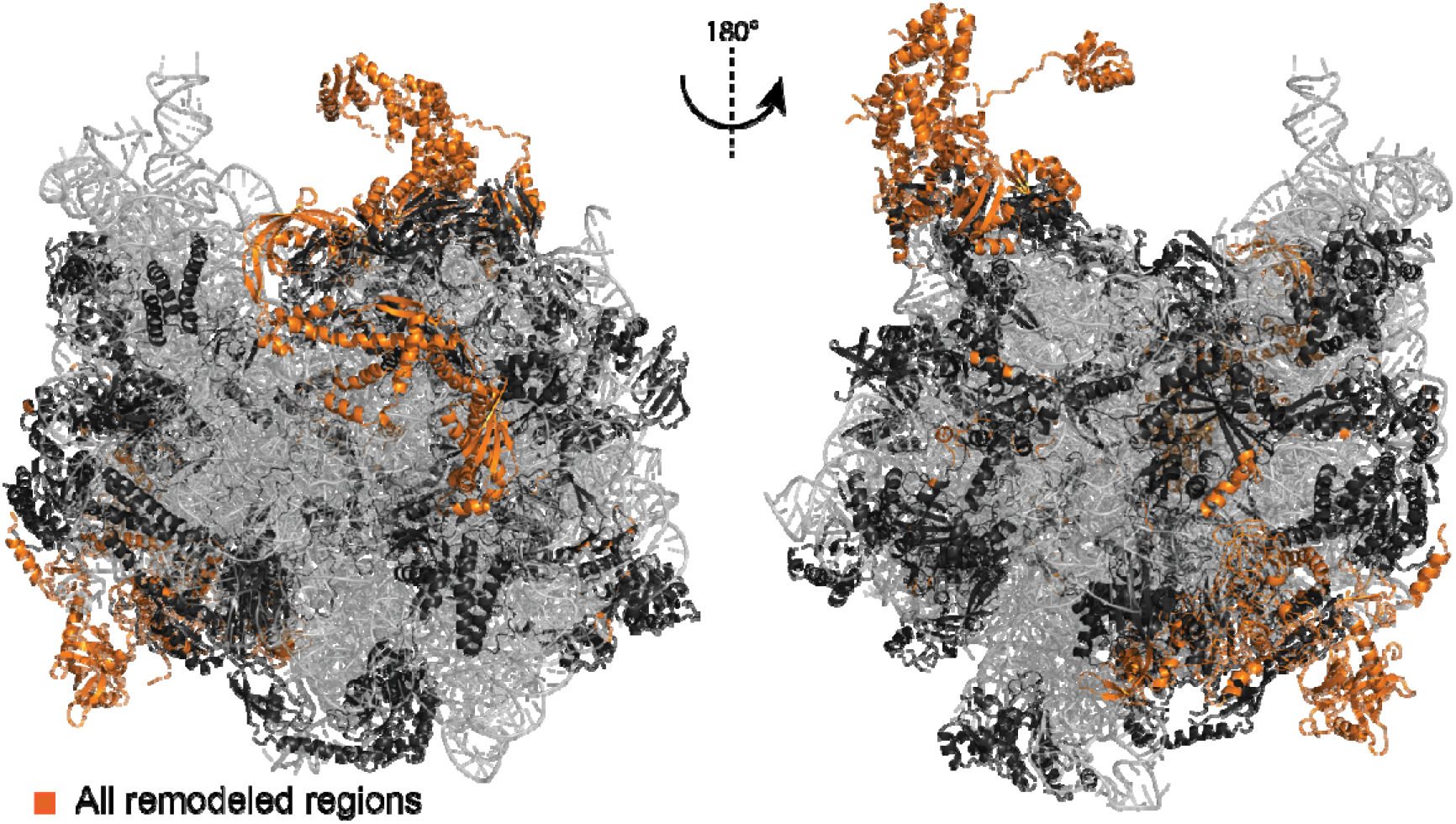
Final model of the *E. coli* 70S ribosome. Ribosomal proteins (black) and rRNA (grey) are shown in cartoon presentation. Added or altered ribosomal protein regions are highlighted in orange.

To conclude, high-resolution methods cannot capture completely the conformational variation and dynamics of large macromolecular complexes and the current work shows that XL-MS is an essential method to identify technical discrepancies, conformational variability, and additional interactors, and ultimately complete and extend the available structural data for the *E. coli* ribosome.

Biomolecules are inherently flexible and highly dynamic, and this underlying flexibility manifests substantially to larger biomolecular structures. As in the case of the ribosome, it is frequently observed in density reconstructions with very high local heterogeneity. These regions are hard to be explicitly modeled, and appear at low resolution. In addition, modeling in low resolution densities often results in a single-state model, ignoring the fact that a lower-resolution density probably reflects structural variation. XL-MS can be highly beneficial for understanding local heterogeneity in unresolved or partially resolved regions and, therefore, deliver complementary information, which may be valuable to ultimately describe conformational variation in biomolecular complexes at high resolution.

## Methods

### Cross-linking

33.3 mg/ml (13.3 μM) *E. coli* ribosome was purchased from New England Biolabs, cat. no. P0763S20. 20 μL of the commercial solution were diluted to 200 μL with 20 mM HEPES (pH 7.6), 10 mM MgAc_2_, 30 mM KCl, and 2 mM TCEP. Four 49-μL aliquots were pipetted into four vials. DSAU was purchased from CFPlus Chemicals, cat. no. PCL006_0050. DSAU was dissolved in neat ACN to a final concentration of 5 mM by sonicating for 5 minutes at room temperature. 1 μL of DSAU stock solution was added to each 49 μL aliquots of 1.33 μM ribosome. The mixtures were incubated for 60 minutes at room temperature. 1 μL of 1 M ammonium bicarbonate was added and incubated for 15 minutes at room temperature for quenching the cross-linking reactions.

### Digestion

50 μl of each cross-linked sample were digested by SMART Digest Trypsin Kit (Thermo Fisher Scientific), by adding 150 μL of SMART Digest buffer containing 5 μg of beads at 70°C for 3 h. The samples were cooled to room temperature and centrifuged for removing the trypsin beads. The samples were reduced by adding 4 mM of DTT and incubating for 30 min at 56°C. 8 mM iodoacetamide was added and incubated 20 min at room temperature in the dark. Alkylation was quenched by adding 4 mM of DTT. Finally, the samples were acidified by adding TFA for LC/MS analysis.

### Mass Spectrometry

The digested samples were analyzed in two technical replicates by LC/MS/MS on an UltiMate 3000 RSLC nano-HPLC system (Thermo Fisher Scientific) coupled to an Orbitrap Q-Exactive Plus mass spectrometer (Thermo Fisher Scientific) equipped with Nanospray Flex ion source (Thermo Fisher Scientific). Peptides were separated on reversed phase C18 columns (precolumn: Acclaim PepMap 100, 300 μm × 5 mm, 5μm, 100 Å (Thermo Fisher Scientific); separation column: packed Picofrit nanospray C18 column, 75 μM × 250 mm, 1.8 μm, 80 Å, tip ID 10 μm (New Objective)). After online desalting, peptides were eluted and separated using a linear water-acetonitrile gradient from 3% to 40% B (with solvent B: 85% (v/v) acetonitrile, 0.1% (v/v) formic acid) over 360 min, 42% to 99% B (5 min) and 99% B (5 min) at a flow rate of 300 nl/min. The separation column was kept at 45°C using an external column heater (Phoenix S&T).

Data were acquired in data-dependent MS/MS mode with stepped higher-energy collision-induced dissociation (HCD) and normalized collision energies of 27%, 30%, and 33%. Each high-resolution full scan (*m/z* 299 to 1799, R = 140,000 at *m/z* 200) in the orbitrap was followed by 10 high-resolution product ion scans (R = 35,000), starting with the most intense signal in the full-scan mass spectrum (isolation window 2 Th); the target value of the automated gain control was set to 3,000,000 (MS) and 250,000 (MS/MS) and maximum accumulation times were set to 100 ms (MS) and 250 ms (MS/MS). Precursor ions with charge states < 3+ and > 7+ were excluded from fragmentation. Dynamic exclusion was enabled (duration 60 seconds, window 2 ppm).

### Data Analysis

Mass spectrometric raw files were converted to mzML using Proteome Discoverer 2.0. Analysis of cross-links was performed with MeroX (version 2.0 beta) using *E. coli* proteins as database (uniprot.org) (FASTA file is included in the Supporting Information). The following settings were applied: Proteolytic cleavage: *C-ter* at Lys and Arg with 3 missed cleavages, peptide length 4 to 30 aa, fixed modification: alkylation of Cys by IAA, variable modification: oxidation of Met, cross-linker: DSBU with specificity towards Lys, Ser, Thr, Tyr, *N-ter* for site 1 and Lys and *N-ter* for site 2, analysis mode: RISEUP mode, maximum missing ions: 2, precursor mass accuracy: 4 ppm, product ion mass accuracy: 8 ppm, signal-to-noise ratio: 2, precursor mass correction activated, prescore cut-off at 10% intensity, FDR cut-off: 1%, and minimum score cut-off: 50. The CRAP database was included for a more stringent FDR estimation. Cross-links identified in the two technical replicates of each digested sample were combined. The MS data have been deposited to the ProteomeXchange Consortium via the PRIDE partner repository with the dataset identifier PXD018935.

### Identification of E. coli ribosomal structures

For the identification of suitable structures, the structural information deposited in the UniProt-database was used. For intermolecular cross-links, the structural information of both interaction partners were cross corelated and the PDB IDs of all structures, in which both proteins are included, were selected. For intramolecular cross-links, no cross-correlation was needed. The structures were then filtered by their annotated resolution, only using structures with a resolution of 3.5 Å or better. NMR structures that do not have a deposited resolution were always included.

### Mapping of cross-links and estimating distance violation

Each identified structure was downloaded from the RCSB protein data bank in the modern mmCIF-format and residue type, residue number, chain identifier, and coordinates of all Cα atoms were manually isolated. This simplified structure file was used for all further analysis. For each cross-link/structure pair, a local sequence alignment for both residues were performed, to guarantee the correct distance measurement and to compensate for annotation discrepancies between the structure and the UniProt annotated sequence (*e.g.* start Met excluded in structure and thereby residue numbering shifted by 1). To perform the alignment, a tripeptide with the cross-linked residue in the middle was fetched from UniProt and aligned to the sequence of the structure, assuming correct residue numbering. For terminal residues, only a dipeptide was used, but the terminal position was considered. If the first alignment failed, the tripeptide was shifted in a +/− 5 residue window around the assumed position in the structural sequence. If the alignment failed (*e.g.* residue not resolved; incorrectly numbered structure), no unambiguous measurement of the distance was possible and the respective cross-link/structure pair were excluded. After successful sequence alignment, the distance *d* between the Cα atoms were calculated with 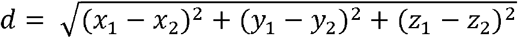, in which P(x_1_, y_1_, z_1_) and P(x_2_, y_2_, z_2_) are the coordinates of both Cα atoms. To differentiate distances in monomeric ribosomal structures from distances in asymmetric units, which would lead to false positive results, the chain identifier label was compared. According to common convention, chains in asymmetric units are distinguishable, and distances from asymmetric units were identified by following rules: (a) first chain uppercase but second chain lowercase or *vis-à-vis* (*e.g.* A to b instead of A to B); (b) if the chain identifier does have two characters, but the first character is differently (*e.g.* AA to BB, instead of AA to AB). All distances were classified by the given threshold of 37.5 Å and categorized in “always satisfied” when all distances of a given cross-link pair are below the threshold, “always violated” when all distances are above the threshold and “mixed-type”, when distances are both, below and above the threshold. Cross-links not mappable to any structure were classified as “novel”.

### Modeling methods

For remodeling, homology modeling, and *de novo* addition of residues MODELLER (Version 9.22) [56] was used as descripted in the manual (chapter 2.2.2), including VTFM and MD optimization of the generated models. The published ribosomal structure “3.2 A cryo-EM ArfA-RF2 ribosome rescue complex (Structure II)” (PDB ID: 5U9F) [29] was used as the starting point for modeling, but the bound ribosome-rescue factor A was removed. Superimposing of the trigger factor was done with PyMOL. To reduce calculation time, only proteins and rRNA within a distance of 10 Å and cross-linked interaction partners of the target protein were considered, and the remodeling was only applied to this single protein, while all others were kept static. Homologue structures for unknown regions in S1 and S3 were identified using HHPRED. The identified structures (PDB ID: 4Q7L for S1 [57]; PDB ID: 2JPL for S3 [41]) were used to model the unknown parts in their ribosomal environment. Missing residues were added in the alignment file and added by *de novo* modeling. Cross-links were included as Gaussian distributed distance restrains (see MODELLER manual chapter 2.2.11). The mean distance of the constraint was stepwise reduced from 45 Å to 25 Å (step size 5 Å) or until the cross-link distance was below the threshold. The standard deviation of the distance restrains was 0.1. The stepwise remodeling was applied to maintain structural integrity of flexible domains and regions. Remodeled proteins were used to replace their incomplete or incorrect counterparts in the working model by using the alignment function from PyMOL, sequentially building an extended ribosomal structure.

## Acknowledgments

The authors would like to thank Mr. Daniele Ubbiali and Ms. Xiaohan Wang for their contribution in XL-MS data analysis. A.S. and P.L.K. acknowledge funding by the Deutsche Forschungsgemeinschaft (DFG, German Research Foundation), RTG 2467, project number 391498659 “Intrinsically Disordered Proteins – Molecular Principles, Cellular Functions, and Diseases”. A.S. acknowledges financial support by the DFG (project Si 867/15-2) and the region of Saxony-Anhalt. P.L.K and C.T. are supported by the Federal Ministry for Education and Research (BMBF, ZIK program, grant number 02Z22HN23 to P.L.K), the European Regional Development Funds for Saxony-Anhalt (grant number EFRE: ZS/2016/04/78115 to P.L.K.) and the Martin Luther University Halle-Wittenberg. C.I. acknowledges funding by the Alexander von Humboldt Foundation and H2020 MSCA IF 2018 EU project 843585.

## Author contributions

C.T. performed analysis and modeling, C.I. peformed cross-linking experiments and data analysis, C.H.I. performed mass spectrometry, A.S. and P.L.K. supervised the study. All authors designed the study, conceived analysis, wrote, and reviewed the manuscript.

## Data availability

Data are available via ProteomeXchange with identifier PXD018935. All models and modeling protocols are available from the corresponding authors upon request.

## Supplementary Information

The Supplementary Information contains the DSAU chemistry, Supplementary Figures and Supplementary Tables as Supplementary Methods, and five Supplementary Tables (Tables S1-S5) as Supplementary Data.

## SUPPLEMENTARY INFORMATION

### Supplementary Methods

#### DSAU chemistry

Considering range in spacer lengths of amine reactive, urea-based MS-cleavable cross-linkers, DSAU stands between DSBU and CDI. DSAU has four carbon atoms less than DSAU and four more than CDI (Scheme 1A). Its C2 spacer arms are the shortest possibility to connect the central MS-cleavable urea moiety (Scheme 1B) and the NHS reactive head groups. DSAU can bridge residues within Cα-C Cα distances of ~25 Å increasing the spatial resolution of DSBU by ~16%, while maintaining the reactivity of all NHS esters-based cross-linkers. Compared to CDI, DSAU is longer, but more selective towards lysine residues. In fact, despite its symmetric structure, CDI is not a homobifunctional cross-linker. Its first imidazole moiety is highly reactive and readily substituted by hydroxy (serines, threonines, and tyrosines) and amine groups (lysines, *N*-termini) in proteins. The resulting intermediate is considerably stable and the second imidazole is slowly displaced almost exclusively by primary amines. The high polarity of the urea moiety, combined with the shorter aliphatic chains of the spacer, makes DSAU less soluble in organic solvents than DSBU and CDI. In our hands, DSAU was solubilized in neat acetonitrile (ACN) up to 5 mM by sonication. This limits the final concentration of DSBU in the reaction mixture. To keep the ACN concentration below 2% (v/v) in the protein buffer, the concentration of DSAU will be 100 μM. More concentrated DSAU stock solutions might be prepared by replacing ACN with hexafluoroisopropanol. Alternatively, suspensions of 20-30 mM DSAU can be pipetted into ACN and then added to the protein mixture.

DSAU was synthesized following the same procedure of DSBU and using glycine as starting material instead of γ-aminobutyric acid (Scheme 1C). DSAU is also commercially available and was purchased from CFPlus Chemicals.

**Scheme 1.**
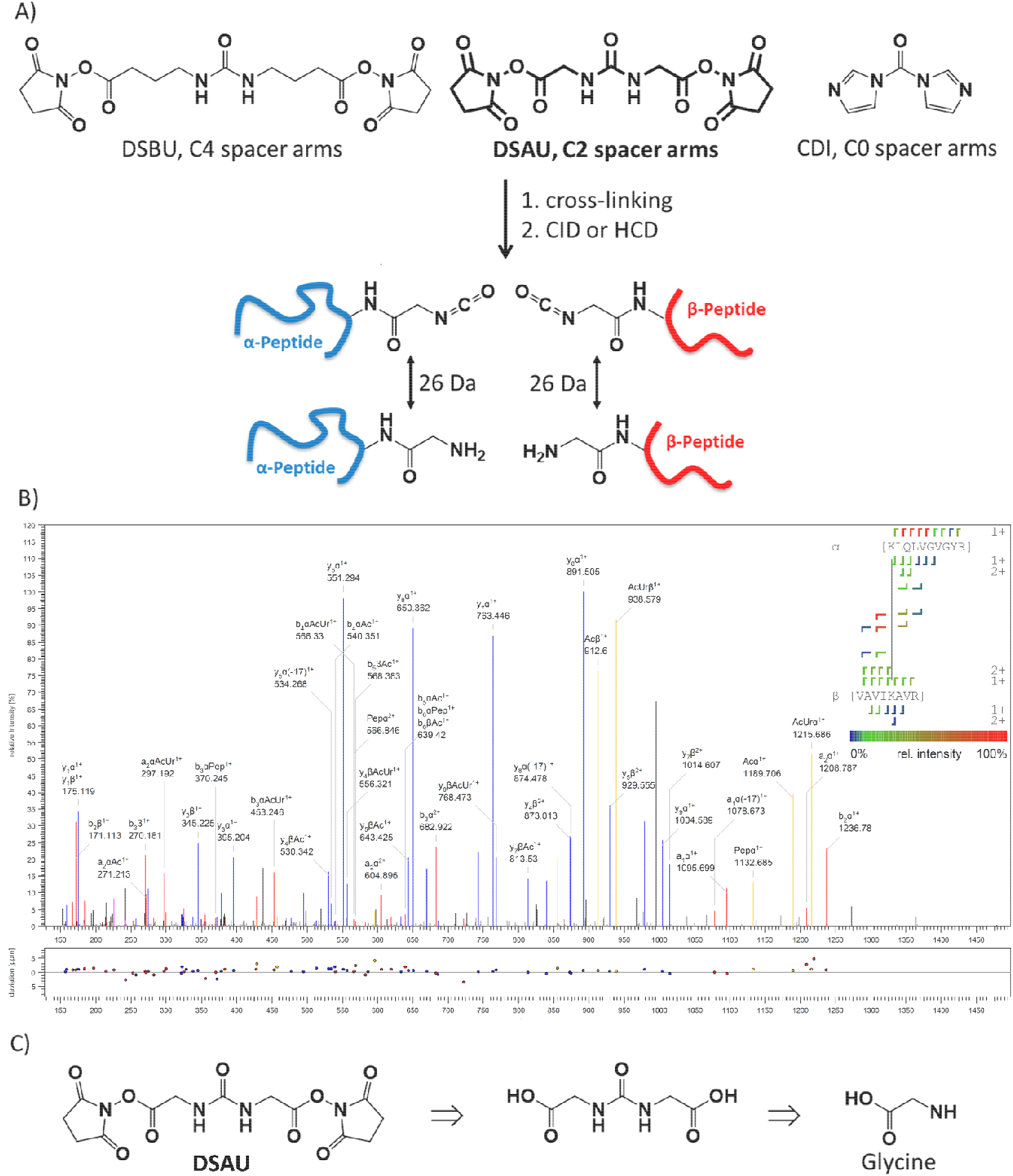
A) Structures of the amine reactive, urea-based MS-cleavable cross-linkers disuccinimidyl dibutyric urea (DSBU), diacetyl dibutyric urea (DSAU) (highlighted in bold), and 1,1’-carbonyldiimidazole (CDI), exhibiting different spacer lengths. B) Fragment ion mass spectrum of a DSAU reaction product, assigned by MeroX 2.0 for a precursor ion at *m/z* 709.764, charge state 3+, corresponding to an intermolecular cross-linked product between K86 of RL6 and K71 of RL7. DSAU reacts mainly with amine groups. Upon fragmentation, DSAU yields two doublets (yellow signals) with a mass difference of 25.979 u in the MS/MS spectrum, facilitating automated data analysis. C) Retrosynthetic pathway of DSAU.

### Supplementary Figures

**Figure S1.**
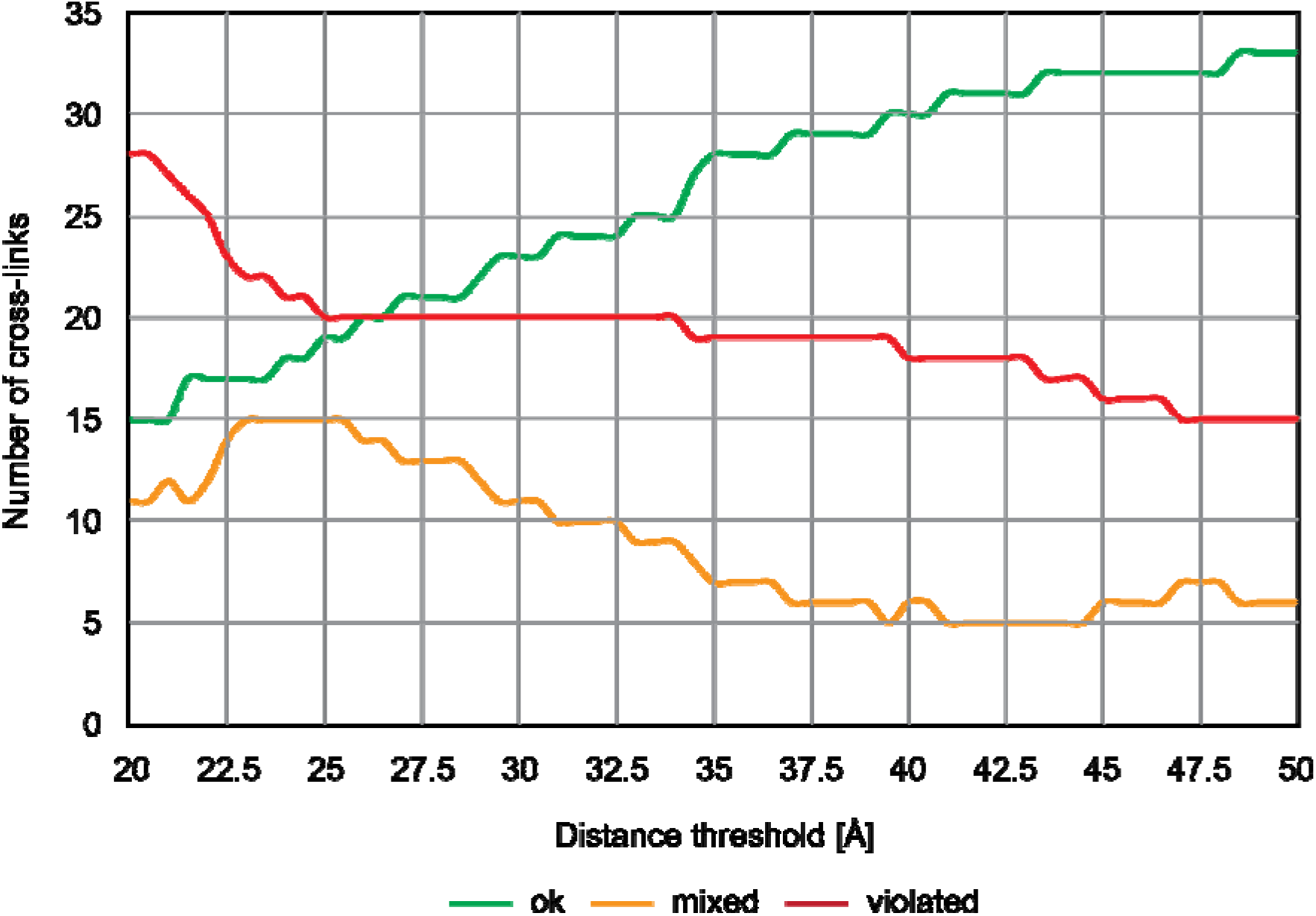
Distribution of cross-link satisfaction and violation using different thresholds. Cross-links are classified as ‘ok’, if they are satisfied for all identified structures and ‘violated’ when they are always above the threshold. ‘Mixed’ cross-links showed both, satisfaction and violation, for different structures indicating flexibility in the respective regions of the *E. coli* ribosome.

**Figure S2.**
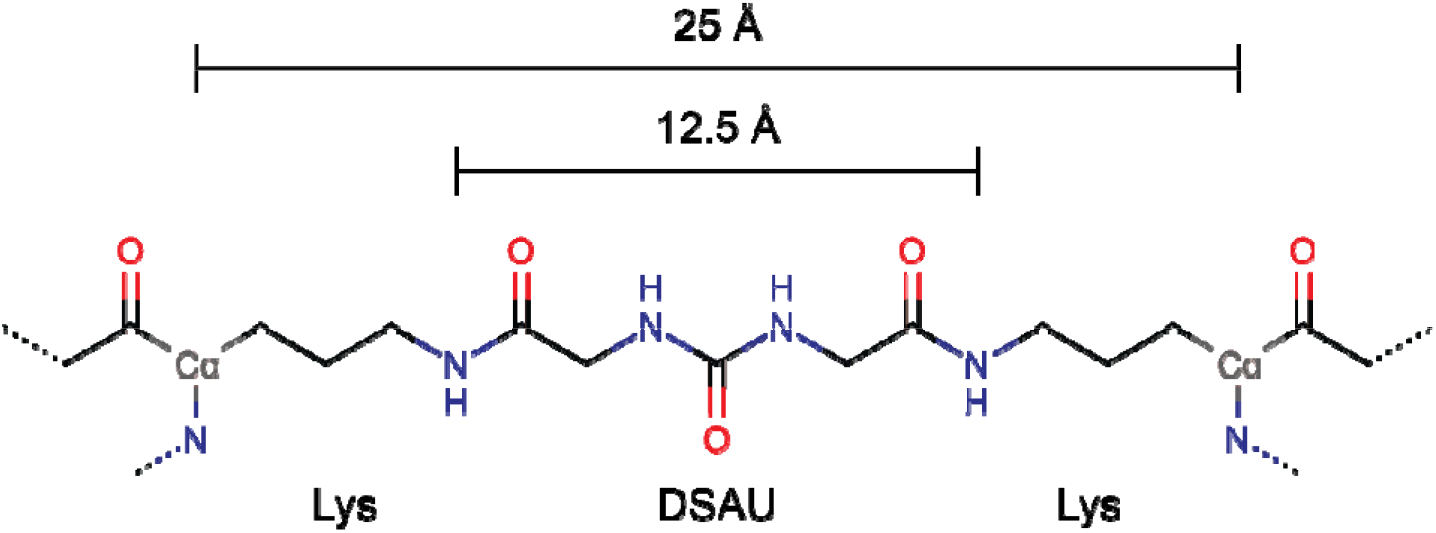
Schematic view of a Lys-Lys cross-link generated by DSAU. The maximum Cα-Cα distance between the connected lysines is 25 Å.

**Figure S3.**
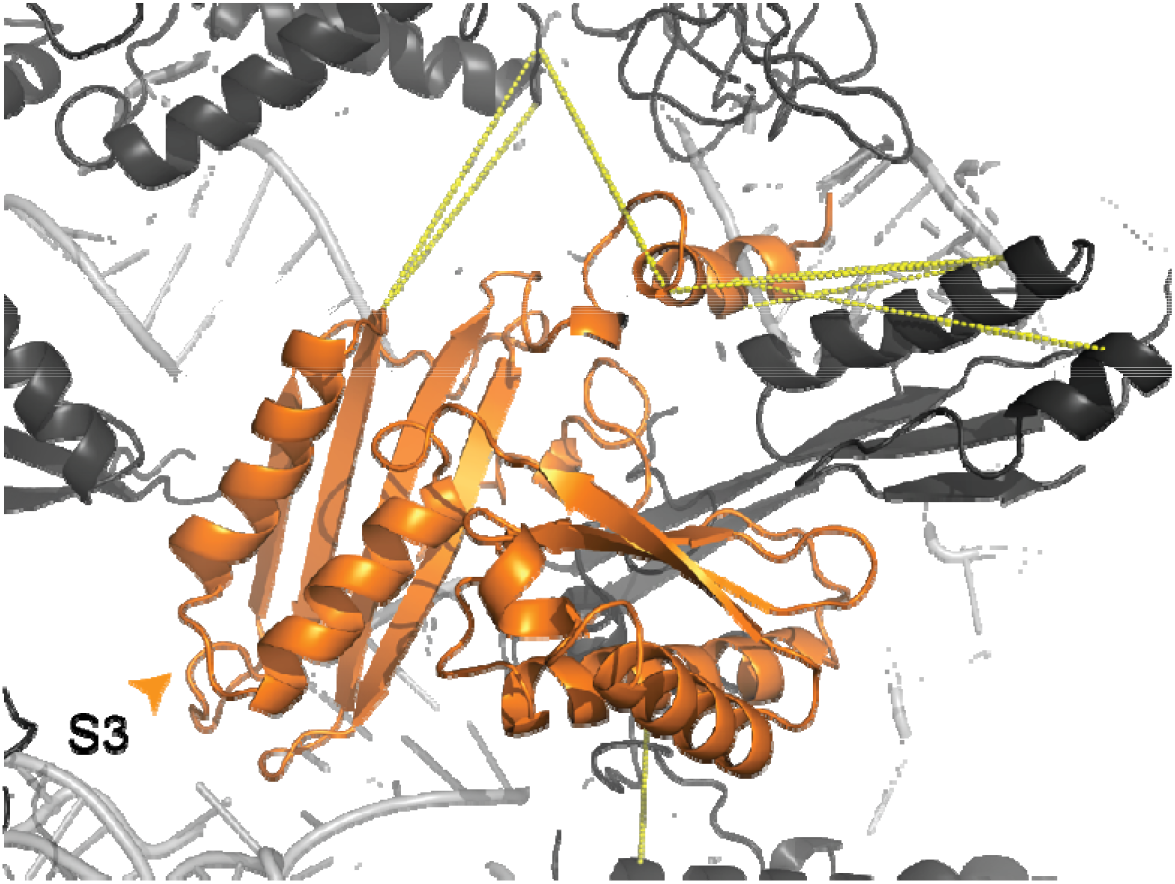
3D Model of S3 satisfying all cross-links without cross-link driven remodeling. (A) Structural model of S3 (orange) after reconstruction of the *C*-terminal region. Ribosomal proteins (black) and rRNA (grey) are shown in cartoon presentation. Cross-links are shown as yellow dotted lines.

**Figure S4.**
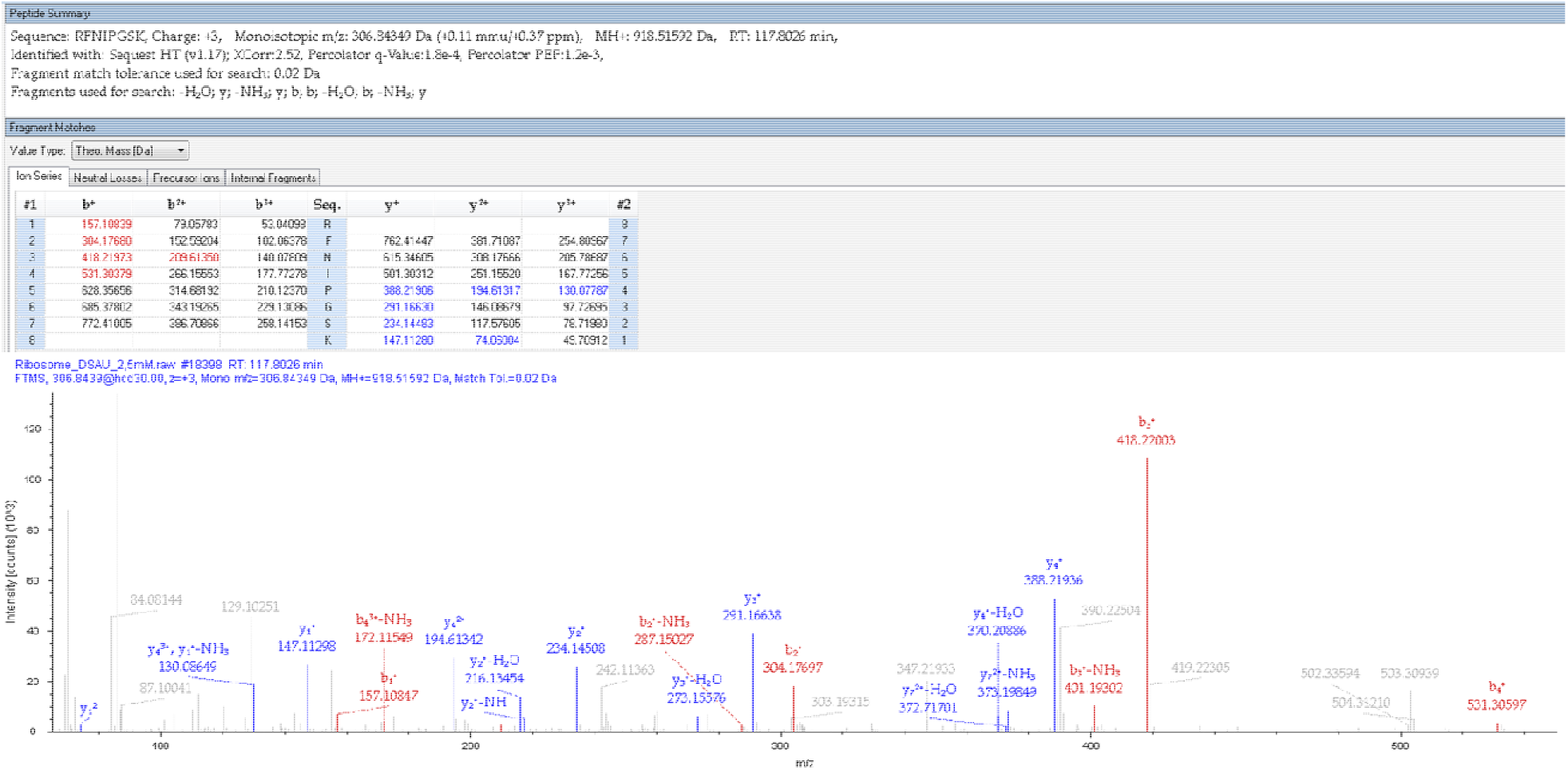
Fragment ion mass spectrum of the C-terminal peptide of L31 (amino acids 62-70, RFNIPGSK). The spectrum was annotated by the Sequest HT algorithm in Proteome Discoverer 2.4.

**Figure S5.**
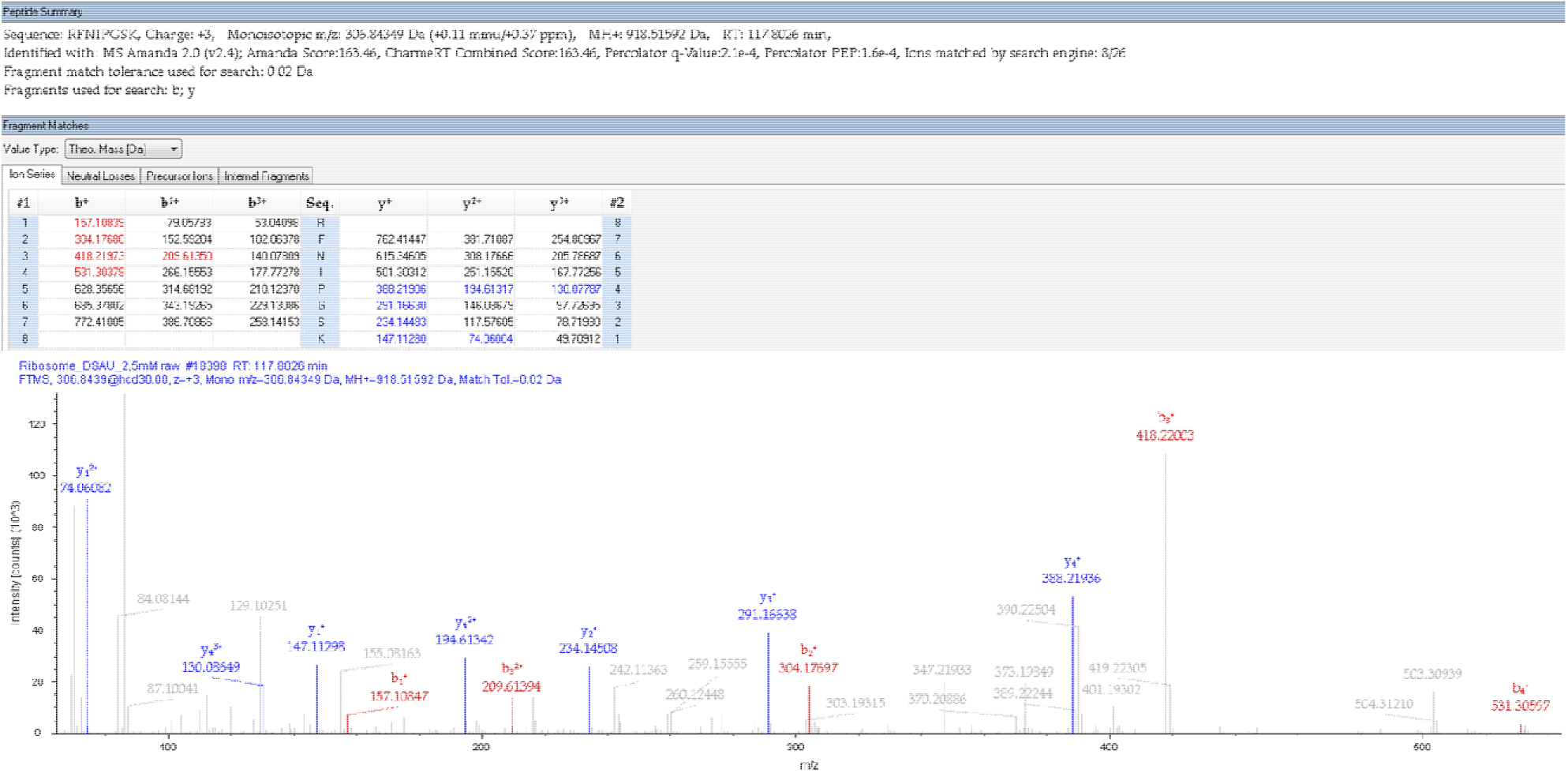
Fragment ion mass spectrum of the C-terminal peptide of L31 (amino acids 62-70, RFNIPGSK). The spectrum was annotated by the Amanda 2.0 algorithm in Proteome Discoverer 2.4.

### Supplementary Tables and Legends

**Table S1. Summary of cross-link pairs.** Intermolecular cross-links are marked with “#”, intramolecular cross-links with “§”. **Table S2. Summary of intramolecular cross-links.** PDB ID, chain identifier, amino acid numbers, the distances (in Å) are given for cross-linked residues.

**Table S3. Summary of intermolecular cross-links.** PDB ID, chain identifier, amino acid numbers, the distances (in Ångstrom) are given for cross-linked residues.

**Table S4. Intra-molecular cross-link mapping of L7/L12.** Cross-links were evaluated in monomeric or multimeric models with Xwalk [1].

**Table S5. Summary of cross-links for the final model.** Cross-link IDs are annotated according to Table S1. Protein name and UniProt Identifier are given. Distances are given in Å and cross-links that could not be mapped are indicated (n.m.). The distances are classified as follows: Satisfied (<37.5 Å for inter-, <30 Å for intramolecular cross-links), minor violated (<50 Å for inter-, <40 Å for intramolecular), and violated. The classification from large-scale mapping is given in brackets.

## References

1. S. Klinge & J. L. Woolford, Jr. Ribosome assembly coming into focus. Nat Rev Mol Cell Biol 20, 116–131, doi:10.1038/s41580-018-0078-y (2019).

2. T. Wagenknecht, J. M. Carazo, M. Radermacher & J. Frank. Three-dimensional reconstruction of the ribosome from Escherichia coli. Biophys J 55, 455–464, doi:10.1016/S0006-3495(89)82839-5 (1989).

3. A. Yonath et al. Characterization of single crystals of the large ribosomal particles from Bacillus stearothermophilus. J Mol Biol 187, 633–636, doi:10.1016/0022-2836(86)90342-6 (1986).

4. H. F. Noller & J. B. Chaires. Functional modification of 16S ribosomal RNA by kethoxal. Proc Natl Acad Sci U S A 69, 3115–3118, doi:10.1073/pnas.69.11.3115 (1972).

5. A. K. Falvey & T. Staehelin. Structure and function of mammalian ribosomes. I. Isolation and characterization of active liver ribosomal subunits. J Mol Biol 53, 1–19, doi:10.1016/0022-2836(70)90042-2 (1970).

6. B. T. Wimberly et al. Structure of the 30S ribosomal subunit. Nature 407, 327–339, doi:10.1038/35030006 (2000).

7. H. E. Huxley & G. Zubay. Electron microscope observations on the structure of microsomal particles from Escherichia coli. J Mol Biol 2, 10–18, doi:10.1016/S0022-2836(60)80003-4 (1960).

8. X. C. Bai, I. S. Fernandez, G. McMullan & S. H. Scheres. Ribosome structures to near-atomic resolution from thirty thousand cryo-EM particles. Elife 2, e00461, doi:10.7554/eLife.00461 (2013).

9. A. Brown & S. Shao. Ribosomes and cryo-EM: a duet. Curr Opin Struct Biol 52, 1–7, doi:10.1016/j.sbi.2018.07.001 (2018).

10. D. Matzov et al. The cryo-EM structure of hibernating 100S ribosome dimer from pathogenic Staphylococcus aureus. Nat Commun 8, 723, doi:10.1038/s41467-017-00753-8 (2017).

11. F. Brandt et al. The native 3D organization of bacterial polysomes. Cell 136, 261–271, doi:10.1016/j.cell.2008.11.016 (2009).

12. R. Kohler, R. A. Mooney, D. J. Mills, R. Landick & P. Cramer. Architecture of a transcribing-translating expressome. Science 356, 194–197, doi:10.1126/science.aal3059 (2017).

13. G. Demo et al. Structure of RNA polymerase bound to ribosomal 30S subunit. Elife 6, doi:10.7554/eLife.28560 (2017).

14. N. Garreau de Loubresse et al. Structural basis for the inhibition of the eukaryotic ribosome. Nature 513, 517–522, doi:10.1038/nature13737 (2014).

15. S. Arenz et al. Structures of the orthosomycin antibiotics avilamycin and evernimicin in complex with the bacterial 70S ribosome. Proc Natl Acad Sci U S A 113, 7527–7532, doi:10.1073/pnas.1604790113 (2016).

16. N. R. Genuth & M. Barna. The Discovery of Ribosome Heterogeneity and Its Implications for Gene Regulation and Organismal Life. Mol Cell 71, 364–374, doi:10.1016/j.molcel.2018.07.018 (2018).

17. F. Liu, D. T. Rijkers, H. Post & A. J. Heck. Proteome-wide profiling of protein assemblies by cross-linking mass spectrometry. Nat Methods 12, 1179–1184, doi:10.1038/nmeth.3603 (2015).

18. P. L. Kastritis et al. Capturing protein communities by structural proteomics in a thermophilic eukaryote. Mol Syst Biol 13, 936, doi:10.15252/msb.20167412 (2017).

19. M. Gotze, C. Iacobucci, C. H. Ihling & A. Sinz. A Simple Cross-Linking/Mass Spectrometry Workflow for Studying System-wide Protein Interactions. Anal Chem 91, 10236–10244, doi:10.1021/acs.analchem.9b02372 (2019).

20. A. Sinz. Divide and conquer: cleavable cross-linkers to study protein conformation and protein-protein interactions. Anal Bioanal Chem 409, 33–44, doi:10.1007/s00216-016-9941-x (2017).

21. C. Iacobucci, M. Gotze & A. Sinz. Cross-linking/mass spectrometry to get a closer view on protein interaction networks. Curr Opin Biotechnol 63, 48–53, doi:10.1016/j.copbio.2019.12.009 (2019).

22. A. Sinz. Chemical cross-linking and mass spectrometry for mapping three-dimensional structures of proteins and protein complexes. J Mass Spectrom 38, 1225–1237, doi:10.1002/jms.559 (2003).

23. R. M. Kaake et al. A new in vivo cross-linking mass spectrometry platform to define protein-protein interactions in living cells. Mol Cell Proteomics 13, 3533–3543, doi:10.1074/mcp.M114.042630 (2014).

24. L. Fischer & J. Rappsilber. Quirks of Error Estimation in Cross-Linking/Mass Spectrometry. Anal Chem 89, 3829–3833, doi:10.1021/acs.analchem.6b03745 (2017).

25. J. Matthew Allen Bullock, J. Schwab, K. Thalassinos & M. Topf. The Importance of Non-accessible Crosslinks and Solvent Accessible Surface Distance in Modeling Proteins with Restraints From Crosslinking Mass Spectrometry. Mol Cell Proteomics 15, 2491–2500, doi:10.1074/mcp.M116.058560 (2016).

26. K. Yugandhar, T.-Y. Wang & H. Yu. Structure-based validation can drastically under-estimate error rate in proteome-wide cross-linking mass spectrometry studies. bioRxiv, 617654, doi:10.1101/617654 (2019).

27. J. Fursch, K. M. Kammer, S. G. Kreft, M. Beck & F. Stengel. Proteome-Wide Structural Probing of Low-Abundant Protein Interactions by Cross-Linking Mass Spectrometry. Anal Chem 92, 4016–4022, doi:10.1021/acs.analchem.9b05559 (2020).

28. M. Q. Muller, F. Dreiocker, C. H. Ihling, M. Schafer & A. Sinz. Cleavable cross-linker for protein structure analysis: reliable identification of cross-linking products by tandem MS. Anal Chem 82, 6958–6968, doi:10.1021/ac101241t (2010).

29. G. Demo et al. Mechanism of ribosome rescue by ArfA and RF2. Elife 6, doi:10.7554/eLife.23687 (2017).

30. M. Diaconu et al. Structural basis for the function of the ribosomal L7/12 stalk in factor binding and GTPase activation. Cell 121, 991–1004, doi:10.1016/j.cell.2005.04.015 (2005).

31. J. Zhou, L. Lancaster, S. Trakhanov & H. F. Noller. Crystal structure of release factor RF3 trapped in the GTP state on a rotated conformation of the ribosome. RNA 18, 230–240, doi:10.1261/rna.031187.111 (2012).

32. A. Pulk & J. H. Cate. Control of ribosomal subunit rotation by elongation factor G. Science 340, 1235970, doi:10.1126/science.1235970 (2013).

33. P. Tian et al. Folding pathway of an Ig domain is conserved on and off the ribosome. Proc Natl Acad Sci U S A 115, E11284–E11293, doi:10.1073/pnas.1810523115 (2018).

34. C. S. Mandava et al. Bacterial ribosome requires multiple L12 dimers for efficient initiation and elongation of protein synthesis involving IF2 and EF-G. Nucleic Acids Res 40, 2054–2064, doi:10.1093/nar/gkr1031 (2012).

35. M. Leijonmarck & A. Liljas. Structure of the C-terminal domain of the ribosomal protein L7/L12 from Escherichia coli at 1.7 A. J Mol Biol 195, 555–579, doi:10.1016/0022-2836(87)90183-5 (1987).

36. A. Kahraman, L. Malmstrom & R. Aebersold. Xwalk: computing and visualizing distances in cross-linking experiments. Bioinformatics 27, 2163–2164, doi:10.1093/bioinformatics/btr348 (2011).

37. B. S. Schuwirth et al. Structural analysis of kasugamycin inhibition of translation. Nat Struct Mol Biol 13, 879–886, doi:10.1038/nsmb1150 (2006).

38. M. Valle et al. Locking and unlocking of ribosomal motions. Cell 114, 123–134, doi:10.1016/s0092-8674(03)00476-8 (2003).

39. L. G. Trabuco et al. The role of L1 stalk-tRNA interaction in the ribosome elongation cycle. J Mol Biol 402, 741–760, doi:10.1016/j.jmb.2010.07.056 (2010).

40. H. Amiri & H. F. Noller. Structural evidence for product stabilization by the ribosomal mRNA helicase. RNA 25, 364–375, doi:10.1261/rna.068965.118 (2019).

41. P. Rogne, G. Fimland, J. Nissen-Meyer & P. E. Kristiansen. Three-dimensional structure of the two peptides that constitute the two-peptide bacteriocin lactococcin G. Biochim Biophys Acta 1784, 543–554, doi:10.1016/j.bbapap.2007.12.002 (2008).

42. L. Zimmermann et al. A Completely Reimplemented MPI Bioinformatics Toolkit with a New HHpred Server at its Core. J Mol Biol 430, 2237–2243, doi:10.1016/j.jmb.2017.12.007 (2018).

43. T. Hesterkamp, S. Hauser, H. Lutcke & B. Bukau. Escherichia coli trigger factor is a prolyl isomerase that associates with nascent polypeptide chains. Proc Natl Acad Sci U S A 93, 4437–4441, doi:10.1073/pnas.93.9.4437 (1996).

44. F. Merz et al. Molecular mechanism and structure of Trigger Factor bound to the translating ribosome. EMBO J 27, 1622–1632, doi:10.1038/emboj.2008.89 (2008).

45. L. Ferbitz et al. Trigger factor in complex with the ribosome forms a molecular cradle for nascent proteins. Nature 431, 590–596, doi:10.1038/nature02899 (2004).

46. B. S. Schuwirth et al. Structures of the bacterial ribosome at 3.5 A resolution. Science 310, 827–834, doi:10.1126/science.1117230 (2005).

47. A. J. Eistetter, P. D. Butler, R. R. Traut & T. G. Fanning. Characterization of Escherichia coli 50S ribosomal protein L31. FEMS Microbiol Lett 180, 345–349, doi:10.1111/j.1574-6968.1999.tb08816.x (1999).

48. M. Duval et al. Escherichia coli ribosomal protein S1 unfolds structured mRNAs onto the ribosome for active translation initiation. PLoS Biol 11, e1001731, doi:10.1371/journal.pbio.1001731 (2013).

49. B. Beckert et al. Structure of a hibernating 100S ribosome reveals an inactive conformation of the ribosomal protein S1. Nat Microbiol 3, 1115–1121, doi:10.1038/s41564-018-0237-0 (2018).

50. A. B. Loveland & A. A. Korostelev. Structural dynamics of protein S1 on the 70S ribosome visualized by ensemble cryo-EM. Methods 137, 55–66, doi:10.1016/j.ymeth.2017.12.004 (2018).

51. A. R. Subramanian. Structure and functions of ribosomal protein S1. Prog Nucleic Acid Res Mol Biol 28, 101–142, doi:10.1016/s0079-6603(08)60085-9 (1983).

52. D. Takeshita, S. Yamashita & K. Tomita. Molecular insights into replication initiation by Qbeta replicase using ribosomal protein S1. Nucleic Acids Res 42, 10809–10822, doi:10.1093/nar/gku745 (2014).

53. P. Salah et al. Probing the relationship between Gram-negative and Gram-positive S1 proteins by sequence analysis. Nucleic Acids Res 37, 5578–5588, doi:10.1093/nar/gkp547 (2009).

54. L. Hoang, K. Fredrick & H. F. Noller. Creating ribosomes with an all-RNA 30S subunit P site. Proc Natl Acad Sci U S A 101, 12439–12443, doi:10.1073/pnas.0405227101 (2004).

55. S. Hong et al. Mechanism of tRNA-mediated +1 ribosomal frameshifting. Proc Natl Acad Sci U S A 115, 11226–11231, doi:10.1073/pnas.1809319115 (2018).

56. A. Sali & T. L. Blundell. Comparative protein modelling by satisfaction of spatial restraints. J Mol Biol 234, 779–815, doi:10.1006/jmbi.1993.1626 (1993).

57. M. A. Bukowska et al. A Transporter Motor Taken Apart: Flexibility in the Nucleotide Binding Domains of a Heterodimeric ABC Exporter. Biochemistry 54, 3086–3099, doi:10.1021/acs.biochem.5b00188 (2015).

## Supplementary References

1. A. Kahraman, L. Malmstrom & R. Aebersold. Xwalk: computing and visualizing distances in cross-linking experiments. Bioinformatics 27, 2163–2164, doi:10.1093/bioinformatics/btr348 (2011).

